# Gnrh1-induced responses are indirect in female medaka Fsh cells

**DOI:** 10.1101/763250

**Authors:** Kjetil Hodne, Romain Fontaine, Eirill Ager-Wick, Finn-Arne Weltzien

**Affiliations:** Department of Basic Sciences and Aquatic Medicine, Faculty of Veterinary Medicine, Norwegian University of Life Sciences, Oslo, Norway

**Keywords:** calcium, cell-cell communication, gnrh receptor, gonadotropins, pituitary

## Abstract

Reproductive function in vertebrates is stimulated by gonadotropin-releasing hormone (GnRH) that controls the synthesis and release of the two pituitary gonadotropins, follicle-stimulating hormone (FSH) and luteinizing hormone (LH). FSH and LH, which regulates different stages of gonadal development, are produced by two different cell types in the fish pituitary, in contrast to mammals and birds, thus allowing the investigation of their differential regulation. In the present work, we show by fluorescent *in situ* hybridization that Lh cells in adult female medaka express Gnrh receptors, whereas Fsh cells do not. This is confirmed by patch clamp recordings and cytosolic Ca^2+^ measurements on dispersed pituitary cells, where Lh cells, but not Fsh cells, respond to Gnrh1 by increased action potential frequencies and cytosolic Ca^2+^ levels. In contrast, both Fsh and Lh cells are able to respond electrically and by elevating the cytosolic Ca^2+^ levels to Gnrh1 in brain-pituitary tissue slices. Using Ca^2+^ uncaging in combination with patch clamp recordings and cytosolic Ca^2+^ measurements, we show that Fsh and Lh cells form homo- and heterotypic networks in the pituitary. Taken together, these results show that the effects of Gnrh1 on Fsh release in adult female medaka is indirect, likely mediated via Lh cells.

## INTRODUCTION

In all vertebrates, it is suggested that gonadotropin-releasing hormone (GnRH) is the main stimulator of the synthesis and release of the two pituitary gonadotropins: follicle-stimulating hormone (FSH) and luteinizing hormone (LH). GnRH neurons originate in the preoptico-hypothalamic area, but whereas in mammals they release their neuropeptide to the portal capillary system at the base of the hypothalamus termed the median eminence (1) and further to the gonadotrope cells through the circulation, teleost fish do not display a typical median eminence. Instead, the neurosecretory fibers from the brain project into the *pars distalis* of the pituitary (2). These neurons either directly innervate the different endocrine cells or terminate in the extravascular space, adjacent to blood capillaries surrounding the endocrine cells (2–5). Additionally, whereas FSH and LH are produced by the same pituitary cells in mammals (6), they are produced by two distinct cell types in teleosts (7–10). This makes teleosts useful models to investigate the differential regulation of FSH and LH synthesis and release.

FSH and LH control different stages of gamete development both in mammals and teleost fish, with FSH mainly stimulating follicular development in females and LH regulating final maturation and ovulation (11–15). In mammals, the differential regulation of FSH and LH appears to depend on a pulsatile release of GnRH, with low-frequency pulses favoring FSH response and high-frequency pulses favoring LH response, especially in terms of gonadotropin subunit gene expression (16–18). Activation of GnRH receptors on gonadotrope cells elicit increased cytosolic Ca^2+^ levels ([Ca^2+^]_i_) with a subsequent alteration in membrane potential and hormone release by exocytosis. A similar response to Gnrh has been shown also in teleost fish (19–25), although a Gnrh pulsatile or frequency dependent release of gonadotropins has not been demonstrated.

As many as 5 or 6 different Gnrh receptor (Gnrhr) paralogs have be identified in several teleost species, as opposed to 1 or 2 paralogs being present in mammals. The high number of Gnrhr paralogs in fish opens the possibility for differential regulation of Fsh and Lh synthesis and release through differential expression of receptors in the two gonadotrope cell types (26–29).

The importance of GnRH signaling seems to differ for the two gonadotropins. In the infertile natural GnRH mutant (*hpg*) mouse, FSH serum levels are reduced by 50%, while LH levels are non-detectable (30, 31). Likewise, in the teleost medaka (*Oryzias latipes*), Gnrh knockout in females prevented ovulation and reduced expression of *lhb*, but did not affect *fshb* expression or follicle development (13). These results suggest that LH cell function is dependent on GnRH stimulation, while adequate FSH synthesis and release may continue in the absence of GnRH. Indeed, some reports suggest that FSH release is more constitutive in nature and closely tied to its synthesis (32), but a detailed model for the differential regulation of FSH and LH is still lacking. In addition to differential regulation by GnRH, precise regulation of FSH and LH release may depend on cell-cell communication. A coordinated GnRH-induced LH release requires communication between LH cells through gap junctions, seemingly in both mammals and teleosts (33, 34).

In this work, we addressed the question, does Gnrh1 regulate Fsh cells and Lh cells through similar direct pathways, or via different mechanisms? To answer this question we used transgenic lines to clearly distinguish Fsh cells from Lh cells, toward the following objectives: 1) Determine whether hypophysiotropic Gnrh1 neurons project to Fsh and/or Lh cells in the medaka pituitary, 2) Determine which *gnrhr* paralog(s) are expressed in medaka Fsh and Lh cells, 3) Characterize the Gnrh1-induced electrical and [Ca^2+^]_i_ responses of medaka Fsh and Lh cells, and 4) Assess capacity for intercellular signaling between gonadotropes in the medaka pituitary.

## MATERIALS AND METHODS

### Animals

In this paper, we used Japanese medaka (*Oryzias latipes*) wild-type (WT, d-rR strain), and four transgenic lines; tg(*gnrh1*:eGfp) (35), tg(*lhb*:hrGfpII) (36), tg(*fshb*:DsRed2) (this paper), double tg(*lhb*:hrGfpII/*fshb*:DsRed2) (this paper). All fish were maintained on a 14:10 hr L:D cycle in a re-circulating system (28°C, pH 7.6 and conductivity of 800 µS) and fed three times a day with either live brine shrimp or pellets (Gemma, Skretting, Stavanger, Norway). Animal experiments were performed according to the recommendations of the Care and Welfare of Research Animals at the Norwegian University of Life Sciences, and under the supervision of authorized investigators.

### Generation of tg(*fshb*:DsRed2) and double tg(*lhb*:hrGfpII/*fshb*:DsRed2) transgenic lines

The medaka *fshb*:DsRed2 transgenic line, tg(*fshb*:DsRed2), was generated using DsRed2-N1 vector (Clontech, California, USA.), digested by NcoI and NotI restriction enzymes (New England Biolabs, USA) and cloned into I-SceI-MCS-leader-Gfp-trailer plasmid, generating a I-SceI-MCS-leader-DsRed2-trailer vector. The 3833 bp endogenous medaka *fshb* promoter sequence was amplified by PCR using primers (see Table 1) with overhang to KpnI and XhoI and cloned into pGEM-T Easy vector. The vector was then amplified, digested with KpnI and XhoI (New England Biolabs, Massachusetts, USA), and the DNA fragment was purified and cloned into the I-SceI-MCS-leader-DsRed2-trailer vector.

**Table 1:**
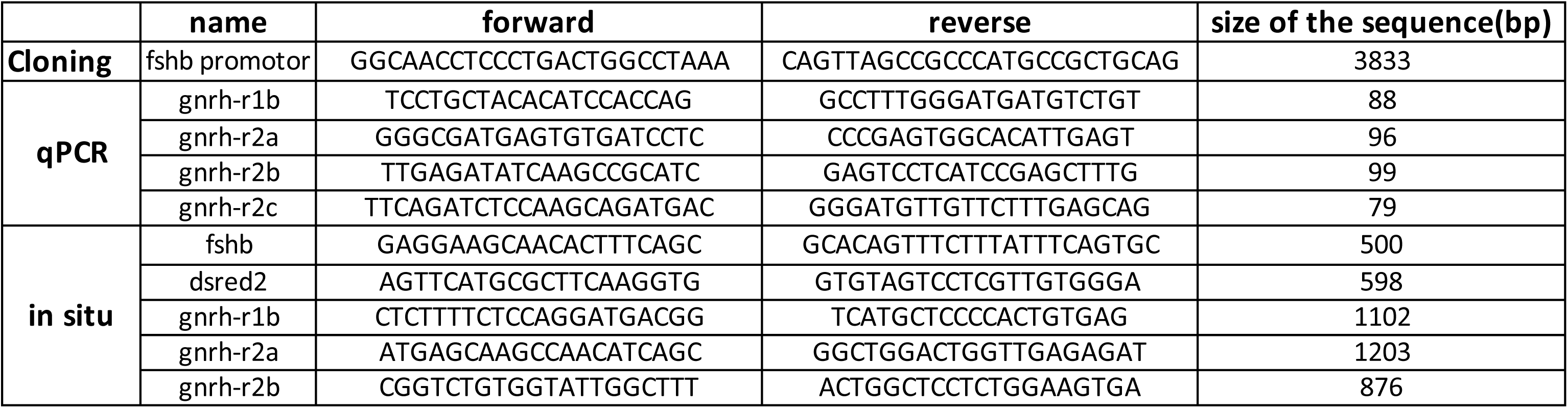
Genes, primers and accession numbers for cloning experiments.

One-cell stage medaka embryos were injected using a manual microinjector (Picospritzer III, Parker Automation, Ohio, USA) with 10 µg/µl transgenic vector diluted in 0.5x commercial meganuclease buffer (Roche Diagnostics, Basel, Switzerland), and with 1 U/µl meganuclease I-SceI and 0.1% phenol red added just prior to use. Injected fish (F0) were raised to adulthood and incrossed. Embryos were screened for DsRed2 by PCR at 5-6 days post fertilization, to determine founders. F0 founders were then crossed with WT and the resulting offspring (F1) were incrossed. Mature F2 were crossed with WT and F2 fish producing 90-100% Rfp-positive progeny were defined to be homozygous: tg(*fshb*:DsRed2). The identified homozygous F2 fish were incrossed to produce a stable homozygous line. Finally, homozygous tg(*fshb*:DsRed2) were crossed with homozygous tg(*lhb*:hrGfpII) to obtain double tg(*lhb*:hrGfp-II/*fshb*:DsRed2) transgenic animals.

### qPCR

Total RNA was isolated from 6 adult female medaka (WT) pituitaries using Trizol (Ambion, USA) and cDNA was prepared from 30 ng total RNA. Specific primers for all target genes (Table 1) were designed with Primer3Plus software and validated based on a series of cDNA dilutions (to assess efficiency and sensitivity) and melting curve analysis (to assess specificity). The qPCR assays were performed as previously described (37) using the LightCycler96 with SYBR Green I (Roche, Switzerland). All qPCR assays were run in duplicate on cDNA diluted 1:5. PCR cycling parameters were 300 s at 95°C followed by 40 cycles at 95°C for 10 s, 60°C for 10 s and 72°C for 6 s, before melting curve analysis to assess qPCR product specificity. Relative expression levels were calculated as described (37), using the combination of three reference genes (*rna18s*, *rpl7*, *gapdh*), according to RefFinder (38).

### Fluorescence *in situ* hybridization

Fluorescence *in situ* hybridization (FISH) for *lhb*, *fshb, dsred2, gnrhr2a, gnrhr1b* and *gnrhr2b* were performed as described previously (39) on 6-month old mature females from tg(*fshb*:DsRed2), tg(*lhb*:hrGfpII) or WT lines (see figure legends for details). First, we investigated the % identity between the RNA probes and the target Gnrhr mRNA (Table 2), and could observed that there were important sequence differences between a RNA probe and the non-targeted *gnrhr* mRNA, thus making unlikely the possibility of unspecific cross reaction Following, riboprobes were cloned into PCR-II TOPO (Thermo Fisher Scientific, Massachusetts, USA) or PGEM-T Easy Vector (Promega, Wisconsin, USA) using primers shown in Table 1 and cDNA from medaka brain and pituitary total RNA. After PCR with M13 reverse and forward primers, sense and antisense probes were synthesized using SP6 or T7 polymerase (Promega) and conjugated with either digoxygenin (DIG, Roche) or fluorescein (FITC, Roche). The fish were euthanized with an overdose of tricaine (MS222; Sigma-Aldrich, Missouri, USA) and tissues fixed by cardiac perfusion with 4% paraformaldehyde (PFA; Electron Microscopy Sciences, Pennsylvania, USA). Free floating 60 μ parasagittal sections of brain and pituitary were made with a vibratome (VT1000S Leica, Wetzlar, Germany). Tissue slices were incubated with hybridization probes for 18 h at 55°C and then incubated with sheep anti-DIG and anti-FITC antibodies conjugated with peroxidase (POD; 1:250; Roche) for probe labeling using the InSituPro VSI robot (Intavis, Germany). Finally, the signal was revealed using custom made TAMRA-conjugated and FITC-conjugated tyramides (39).

**Table 2:** Percent sequence identity between *gnrhr* RNA probes and the mRNA for the different *gnrhr* paralogs in medaka.

Percentage identity
| probe/mRNA | gnrh-r1b | gnrh-r2a | gnrh-r2b |
| --- | --- | --- | --- |
| gnrh-r1b | 100 | 56.2 | 57.7 |
| gnrh-r2a | 56.2 | 100 | 58.2 |
| gnrh-r2b | 41.1 | 41.2 | 100 |

### Immunofluorescence

Immunofluorescence (IF) was performed on free floating 60 μm parasagittal sections as described previously (39). Briefly, brains and pituitaries from 6-month old tg(*gnrh1*:eGfp) females were collected and fixed with 4% PFA. Tissues were then embedded in 3% agarose before sectioning with a vibratome. Then, tissues were blocked for 1 h and incubated with primary custom-made polyclonal rabbit anti-medakaLhβ (1:2000; AB_2732044) and anti-medakaFshβ (1:500; AB_2732042), previously validated (40). For Fshβ IF, epitope retrieval treatment using 2 N HCL (dissolved in phosphate buffered saline solution with 0.1% Tween, PBST) for 1 h at 37°C was necessary prior to blocking and antibody incubation. Signal was amplified with secondary antibody AlexaFluor 555 (1:1000; AB_2535850 Invitrogen). Sections were mounted in Vectashield Antifade Mounting Medium (Vector laboratories, California, USA) and imaged as described below.

### DiI injection

To label pituitary blood vessels, 6-month old siblings of the cross between tg(*gnrh1*:eGfp) and tg(*fshb*:DsRed2), i.e. tg(*gnrh1*:eGfp/*fshb*:DsRed2), were euthanized with an overdose of tricaine. Cardiac perfusion with DiIC18(5)-DS (1,1’-Dioctadecyl-3,3,3’,3’-Tetramethylindodicarbocyanine-5,5’-Disulfonic Acid; Thermo Fisher Scientific) diluted in 4% paraformaldehyde (PFA) was performed as described in (41).

### Confocal Imaging

For whole pituitary imaging, pituitaries from tg(*gnrh1*:eGfp) females were collected, fixed in 4% PFA and mounted between slide and coverslip with spacers in Vectashield Antifade Mounting Medium. All confocal images were acquired using LSM710 confocal microscope (Zeiss, Oberkochen, Germany) with 25X (N.A. 0.8) or 40X (N.A. 1.2) objectives. Channels were acquired sequentially to avoid signal crossover between different filters. Images were processed using ZEN software (version 2009, Zeiss). Z-projections from confocal stacks of images were obtained using Fiji (version 2.0.0 (42)). 3D reconstruction was built using Fiji 3D viewer plugin.

### Primary dispersed pituitary cell culture

Pituitaries were collected from 10 mature, gravid females from the tg(*lhb*:hrGfpII) and tg(*fshb*:DsRed2) lines separately following decapitation and dissociated according to (43). In brief, pooled pituitaries were digested with trypsin (2 mg/mL, Sigma-Aldrich) for 30 min at 26°C, then incubated with trypsin inhibitor (1 mg/mL, Sigma-Aldrich) and DNase I Type IV (2 μg/mL, Sigma-Aldrich) for 20 min at 26°C with gentle shaking. Pituitary cells were mechanically dissociated using a glass pipette, centrifuged at 200 *g* and resuspended in growth medium (L-15, Life Technologies) adjusted to 280-290 mOsm with mannitol and to pH 7.75 with 10 mM NaHCO_3_, 1.8 mM glucose, and penicillin/streptomycin (50 U/mL of medium, Lonza, Verviers, Belgium). Dissociated cells were plated on poly-L-lysine pre-coated dishes fitted with a central glass bottom (MatTek Corporation, Ashland, MA, USA). The cell density was low to prevent intercellular contact and cultures were used within 48 h after plating to minimize changes in gene expression. Three different cell cultures were made for patch clamping and for Ca^2+^ imaging. In each culture 2-7 primary target cells were stimulated by 1 μM Gnrh1 (Bachem Americas, Inc. CA, USA).

### Live brain-pituitary slices

Using mature females from the tg(*lhb*:hrGfpII), tg(*fshb*:DsRed2), and tg(*lhb*:hrGfpII/*fshb*:DsRed2) lines, 150 μm brain-pituitary sections were made according to (44). In brief, following decapitation, the brain-pituitary complex was removed and embedded in 2% agarose (Sigma-Aldrich) dissolved in Ca^2+^- and BSA-free Extracellular Solution (ECS) (mM): NaCl 134, KCl 2.9, MgCl_2_ 1.2, HEPES 10, and glucose 4.5. The solution was adjusted to pH 7.75 with 1 M NaOH and osmolality adjusted to 290 mOsm with mannitol before filter sterilization. Following sectioning, the brain-pituitary slices were moved to the recording chamber containing ECS with 2 mM Ca^2+^ and 0.1% BSA.

### Electrophysiology

All electrophysiological experiments were conducted using the perforated patch clamp technique in current clamp on either brain-pituitary slices (44) or primary dissociated cells (43) from the tg lines described above (*Live brain-pituitary slices*), and with ECS containing 2 mM Ca^2+^ and 0.1% BSA. The experiments were conducted between 14:00 h and 19:00 h. Patch pipettes were made from thick-walled borosilicate glass with 4-5 MΩ resistance. The following intracellular (IC) electrode solution was added to the patch pipette (mM): KOH 120, KCl 20, HEPES 10, sucrose 20, EGTA 0.2. The pH was adjusted to 7.2 with C_6_H_13_NO_4_S, and osmolality to 280 mOsm with sucrose. To perforate the cell membrane amphotericin B (Sigma-Aldrich) was added to 0.24 mg/ml (see details in (44)). The electrode was coupled to a Multiclamp 700B amplifier (Molecular Devices, California, USA). The recorded signals were digitized at 4 to 10 kHz and filtered at one third of the sampling rate using a Digidata 1550B plus with hum silencer (Molecular Devices). All commands and recorded signals were handled using pClamp 10 software (Molecular Devices). Gnrh1 (Bachem) was dissolved to 1 μM in filtered ECS with 0.1% BSA and applied to brain-pituitary slices or dispersed cells using 20 kPa puff ejection through a 2 MΩ pipette, 30-40 μm from the target cell. The cells were visualized using an Infrared Dot Gradient Contrast (DGC) system coupled to an up-right microscope (Slicescope, Scientifica, Uckfield, UK) with a 40X water immersion objective (N.A. 0.8, Olympus, Shinjuku, Japan). The genetically labeled gonadotrope cells were visualized using a light-emitting diode light source (pE-4000 CoolLED, Andover, UK). DsRed2 was excited at 550 nm and emission collected after passing a 630/75 nm bandpass filter (Chroma, Vermont, USA); hrGFPII was excited at 470 nm and emission was collected after passing a 525/50 nm bandpass filter (Chroma). Data were analyzed using AxoGraph version X 1.7 (www.axograph.com) and Matlab version R2018a (Mathworks, Massachusetts, USA).

### Microfluorimetry

Ca^2+^ imaging on adult female pituitaries was performed separately for tg(*fshb*:DsRed2) and tg(*lhb*:hrGfpII) using the calcium-indicator dyes Cal-590-AM (AAT Bioquest, California, USA) for tg(*lhb*:hrGfpII), and Fluo4-AM (ThermoFisher Scientific, Massachusetts, USA) for tg(*fshb*:DsRed2). Dyes were loaded at 5 μM 30 min (cell culture) or 60 min (brain-pituitary slices), at 27°C in 2 mL BSA-free ECS with 1 μL 20% Pluronic (Sigma), followed by 20 min in ECS with 0.1% BSA to de-esterify the dyes. Cal590 was excited at 580 nm and emission images were collected following passage through a 630 nm wavelength / 75 nm bandwidth filter (ET630/75 nm emitter, Chroma). Fluo4 was excited at 470 nm and emission images were collected following passage through a 525 nm wavelength / 50 nm bandwidth filter (ET525/50 nm emitter, Chroma). Cells were imaged using a sCMOS camera (optiMOS, QImaging, British Columbia, Canada) with 50 to 80 ms exposure and 1-2 Hz sampling frequency. Both light source and camera were controlled by μManger software, version 1.4 (45). Relative fluorescence intensity was calculated after background subtraction as changes in fluorescence (F) divided by the average intensity of the first 15 frames (F_0_). Data were analyzed using Fiji (46).

### Uncaging

Uncaging experiments were performed on tg(*fshb*:DsRed2) and tg(*lhb*:hrGfpII), as well as on the double tg(*lhb*:hrGfpII/*fshb*:DsRed2) line. In these experiments, 5 μM NP-EGTA (caging compound) was loaded into the cells (same procedure as the dye loading described above under Microfluorimetry) to chelate Ca^2+^, followed by uncaging and destruction of the NP-EGTA with 50-250 ms pulses from a 405 nm laser (Laser Applied Stimulation and Uncaging system, Scientifica). The laser was pre-set at 7 mW and passed through an 80/20 beam splitter before targeting the cells. The output effect was not calculated. The laser was guided and targeted at distinct regions of the cells using two galvanometer scan mirrors (one for each axis) controlled by Scientifica software developed in LabVIEW (National Instruments, Texas, USA). Laser reflection artifacts were removed post recording from the final imaging profile.

### General figure making

Composites were assembled using Adobe Photoshop and Indesign CC (Adobe Inc, California, USA). Images were processed using ImageJ open source software (versions 1.37v and 2.0.0, National Institute of Health, USA). Photo montages were made in Adobe Photoshop, Adobe Illustrator, and Adobe Indesign CC2018 (Adobe Inc).

## RESULTS

### Development of new medaka transgenic lines and mapping of gonadotrope cells in medaka pituitary

To study medaka Fsh cells, we developed a new transgenic line (tg(*fshb*:DsRed2)) in which DsRed2 expression is controlled by the cloned 3833 bp endogenous *fshb* promotor. Following confirmation of transgene homozygosity, specificity of the DsRed2 reporter was verified using *in situ* hybridization. Multicolor fluorescent *in situ* hybridization with specific probes for *fshb* and *dsred2* showed nearly complete co-localization of the two probes, confirming that DsRed2 is a reliable marker for *fshb* cells in this line (Figure 1A-D). Fsh cells were located in the median part of the *proximalis pars distalis* (PPD). Homozygous tg(*fshb*:DsRed2) fish were crossed with the previously established tg(*lhb*:hrGfpII) line (Figure 1E-H) established by Hildahl et al (36) and recently validated by Fontaine et al (47), to obtain the double tg(*lhb*:hrGfpII/*fshb*:DsRed2). In the double tg line, the two cell types expressing either *fshb* or *lhb* were clearly separated (Figure 1I-L). Fsh and Lh cells were localized in the PPD, with Fsh cells in close proximity of the Lh cells. However, while Lh cells were localized in the ventral and lateral part of the PPD, Fsh cells were localized more medially.

**Figure 1:**
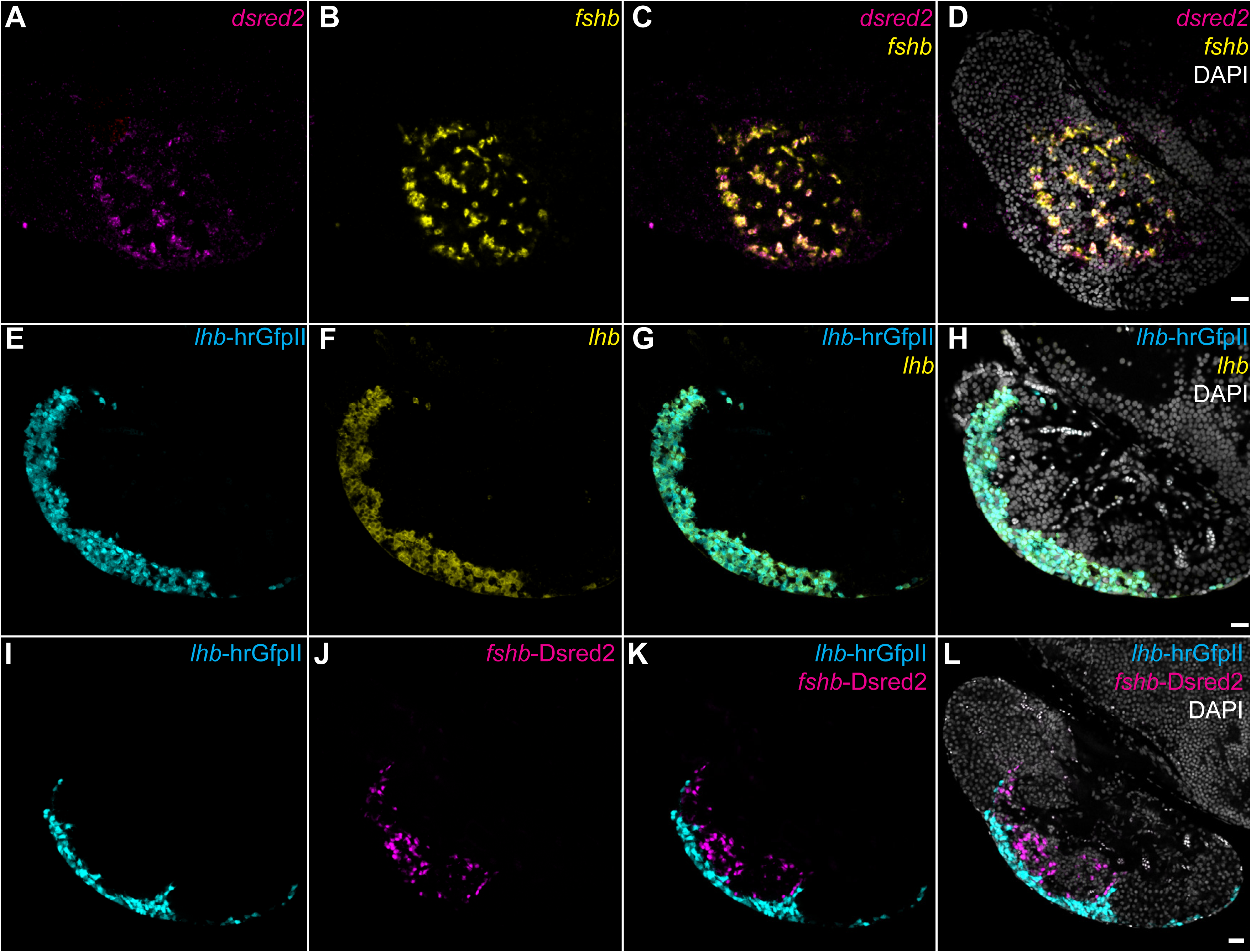
Confirmation of the Fshβ and Lhβ reporter gene expression and generation of the double transgenic line. (A-D) Confocal images of parasagittal sections of medaka brain and pituitary from the *fshb*-DsRed2 transgenic line, (E-H) the *lhb*-hrGfpII transgenic line, or (I-L) the double transgenic line. (A-D) Tissue sections were labeled for *fshb* and *dsred2* by FISH. (E-H) Tissue sections were labeled for *lhb* by FISH. (D), (H) and (L), represent the merged images from the respective left panels together with nuclear (DAPI) staining. Anterior to the left and dorsal to the top. Scale bars: 20 μm.

### Localization of Gnrh1 fibers within the medaka pituitary

To investigate the location of the hypophysiotropic Gnrh1 neuron fibers, we utilized the established tg(*gnrh1*:eGfp) medaka (36). The majority of the PPD was well innervated by Gnrh1 projections (Figure 2A and B), with most fibers reaching the ventral and lateral PPD where Lh cells are located. Immunofluorescence to localize Fshβ (Figure 2C-F) and Lhβ (Figure 2G-J) on tg(*gnrh1*:eGfp) pituitary slices showed that Gnrh1 projections were found in the proximity of both gonadotrope cell types. In fact, Gnrh1 fibers project throughout the whole PPD, passing next to Fsh cells and reaching the ventral and lateral surface, where Lh cells are localized. Using fish perfused with DiI to visualize blood vessels we observed that Gnrh1 projections also followed the path of the blood vessels. (Figure 2K and L).

**Figure 2:**
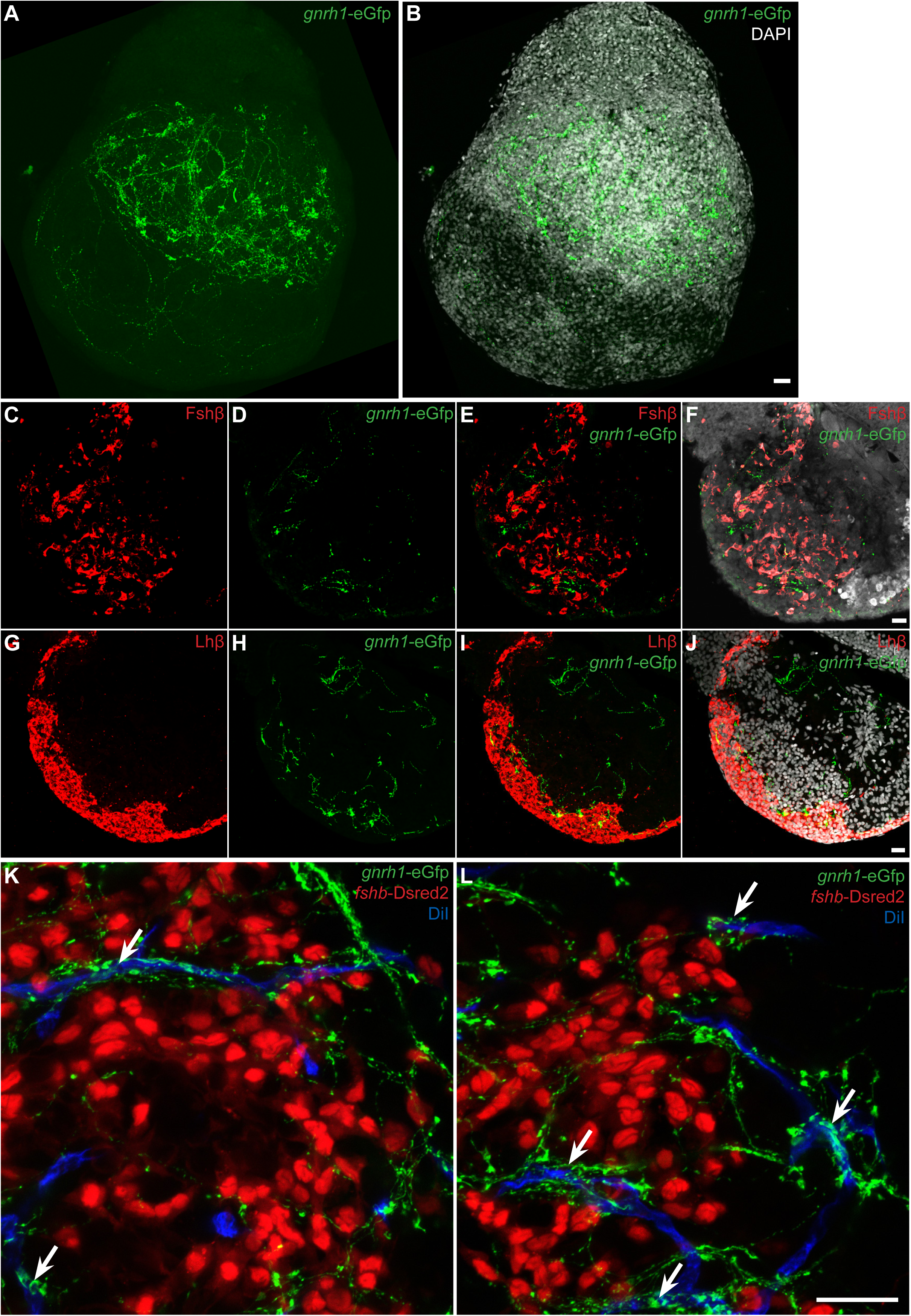
Pituitary innervation by genetically labeled Gnrh1 neurons. (A-B) Z-projections of confocal images from the whole pituitary without (A) or with nuclear (DAPI) staining (B) from the tg(*gnrh1*-eGfp) line. Anterior to the top. (C-J) Confocal images of parasagittal brain-pituitary sections from the tg(*gnrh1*-eGfp) labeled for Fshβ (C-I) or Lhβ (G-J) by IF. (F) and (J) represent the merged images from the respective left panels together with nuclear (DAPI) staining. (K-H) Z-projections (5 μm z-stack) from confocal images of parasagittal brain-pituitary sections from the siblings of the cross between tg(*gnrh1*-eGfp) and tg(*fshb*-DsRed2), i.e. tg(*gnrh1*-eGfp/*fshb*-DsRed2), cardiac perfused with DiI. Scale bars: 20 μm.

### Expression of *gnrhr* in Fsh and Lh cells

To further explore how Gnrh1 regulates Fsh and Lh we used qPCR to measure transcripts of all four *gnrhr* paralogs identified in the medaka genome. While the four genes were cloned from brain RNA, the qPCR assays revealed expression of only three receptor genes in the pituitary (Figure 3A): *gnrhr1b*, *gnrhr2a* and *gnrhr2b*. Expression of *gnrhr2c* was not detected in any of the pituitary samples. To identify cell specific expression, we used double color fluorescent *in situ* hybridization combining each *gnrhr* with either *fshb* or *lhb* (figure 3B-Z). *In situ* probes were designed to minimize the chances for cross reaction between the probes and target *gnrhr* mRNA (Table 2). The results showed that *gnrh1b* is expressed in the posterior part of the pituitary in some of the most posterior *lhb* cells but not in *fshb* cells. *gnrhr2a* is expressed in the ventral surface of the pituitary, almost exclusively in *lhb* cells and never in *fshb* cells. Indeed, a near full co-localization was observed with only a few extra cells expressing *gnrhr2a* without *lhb*. Finally, *gnrhr2b* is expressed in the posterior part of the pituitary, in a similar region to *gnrhr1b*, but was found in very few *lhb* cells (fewer than for *gnrhr1b*), and never in *fshb* cells. Thus, using *in situ* hybridization, we found no evidence of *gnrhr* expression in *fshb* cells.

**Figure 3:**
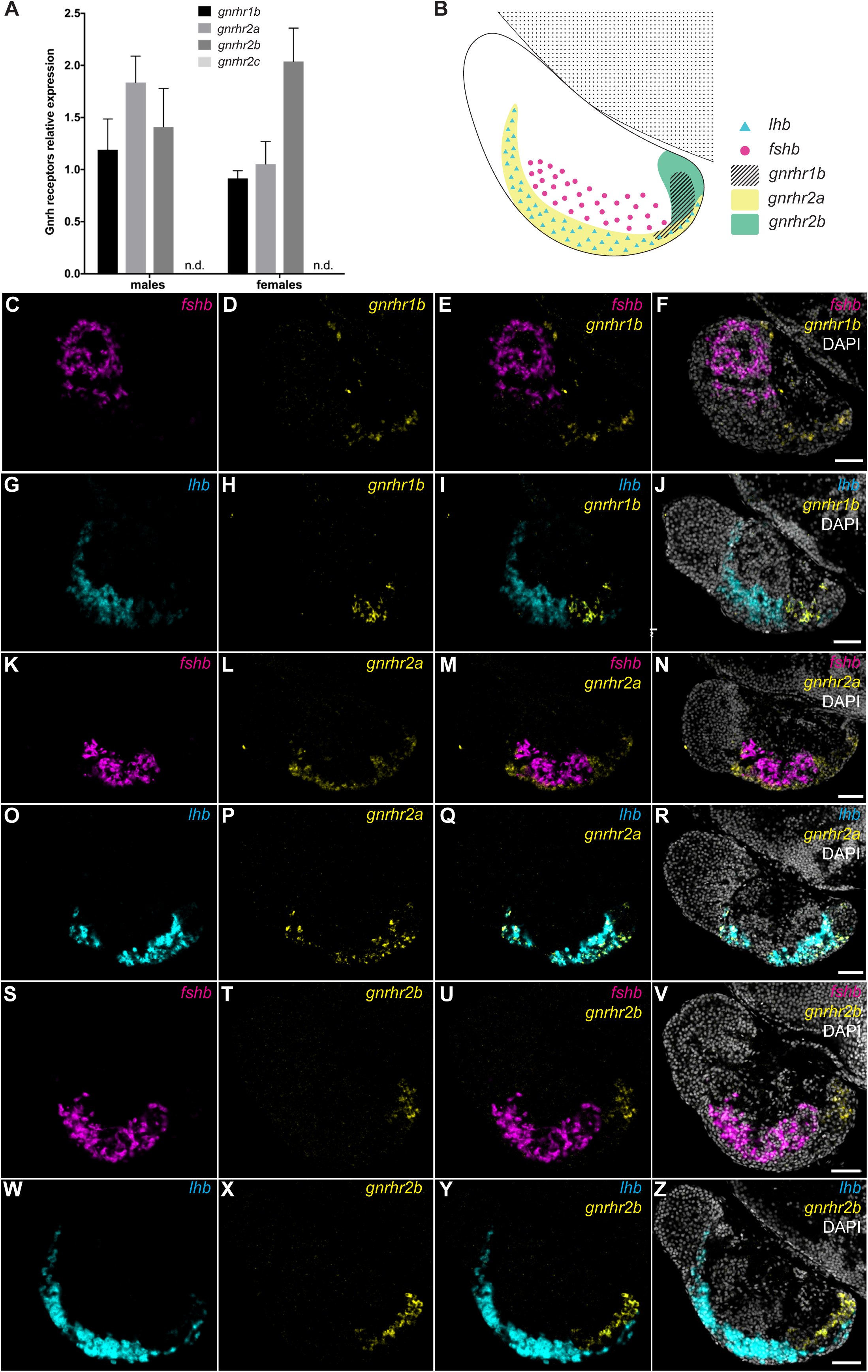
*gnrh* receptor (*gnrhr*) expression in the pituitary. (A) Graphic presenting the relative expression (mean + SEM) of the four *gnrhr* paralogs in both male and female medaka pituitary. (B) Schema presenting the area of expression of the three *gnrhr* expressed in the pituitary and the location of the Fsh and Lh cells therefore summarizing the observations in the remaining panels. (C-Z) Confocal images of brain-pituitary parasagittal sections double-labelled by FISH for *gnrhr1b* together with *fshb* (C-F) or *lhb* (G-J), *gnrhr2a* together with *fshb* (K-N) or *lhb* (O-R) or *gnrhr2b* together with *fshb* (S-V) or *lhb* (W-Z). (F), (J), (N), (R) and (V) represent the merged images from the respective left panels together with nuclear (DAPI) staining. Anterior to the left and dorsal to the top. Scale bars: 50 μm.

### Effect of Gnrh1 on Fsh and Lh cells in dispersed pituitary cell culture

To further explore and validate the *in situ* hybridization results we performed a series of patch clamp and Ca^2+^ imaging experiments in which we stimulated dissociated medaka pituitary cells with Gnrh1. These experiments were conducted 24-48 h after plating the cells.

During the initial current clamp recordings, we observed spontaneous firing of action potentials in 60% (n = 20) of the Fsh cells (Figure 4A), similar to our previous results on medaka Lh cells (21,22,43). The spontaneous firing of Fsh cells occurred in cells with oscillating membrane potential around −45 to −40 mV with the action potential often overshooting 0 mV (liquid junction potential not corrected for). The firing frequency of Fsh cells ranged from 0.5 to 3 Hz.

**Figure 4:**
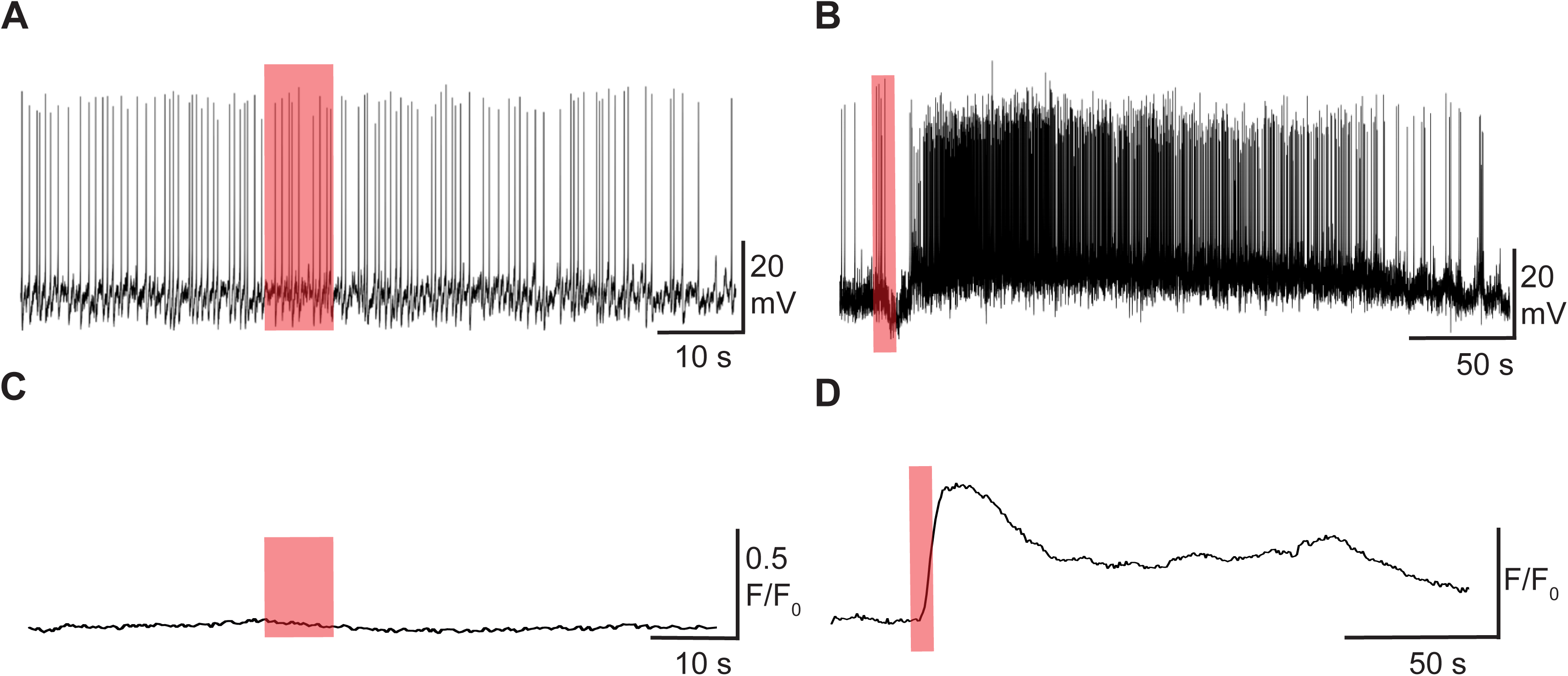
Gnrh1-induced electrical and Ca^2+^ responses of Fsh and Lh cells from primary dispersed cell cultures. The experiments were conducted in dispersed pituitary cells from adult female medaka, using tg(*lhb*:hrGfpII/*fshb*:DsRed2) for electrophysiology and tg(*lhb*:hrGfpII) or (*fshb*:DsRed2) for Ca^2+^ imaging. (A) Typical current clamp recording of a spontaneously firing Fsh cell. Following Gnrh1 application (orange transparent bar overlaying the trace), no apparent changes in the electrical response could be observed (a total of n = 8 cells from 3 different cell cultures combined). (B) Typical current clamp recording of a spontaneously firing Lh cell. Gnrh1 application using puff ejections induced a biphasic response with an initial hyperpolarization followed by depolarization and increased firing and/or increase action potential duration (see also (43)). (C) Ca^2+^ imaging of an Fsh cell. Following Gnrh1 application no response was observed in Fsh cells (a total of n = 10 cells from 3 different cell cultures combined). (D) Gnrh1-induced Ca^2+^ response of an Lh cell. The Lh cells responded to Gnrh1 with a biphasic increase in [Ca^2+^]_i_ mirroring the electrical response in (A) (see details in (21, 22)).

Consistent with the *in situ* hybridization results indicating that Lh cells express *gnrhr* whereas Fsh cells do not, we found a biphasic response (electrical and [Ca^2+^]_i_) to Gnrh1 in Lh cells (Figure 4B and C). The electrical response consisted of an initial membrane hyperpolarization followed by depolarization and a robust increase in action potential frequency (Figure 4B). This response reflects the changes in [Ca^2+^]_i_ with the initial membrane hyperpolarization being due to Ca^2+^ release from internal stores and subsequent activation of Ca^2+^-activated K^+^ channels. This initial release of Ca^2+^ from internal stores is followed by a plateau reflecting the influx of extracellular Ca^2+^ as a result of increased firing frequency (Figure 4D). In contrast to the biphasic response to Gnrh1 observed in Lh cells, we were not able to detect any changes (electrical or [Ca^2+^]_i_) in Fsh cells following Gnrh1 exposure (Figure 4A and C).

### Effect of Gnrh1 on Fsh and Lh cells in live brain-pituitary tissue slices

To further investigate the results observed in cell culture, we utilized brain-pituitary tissue slices from the double tg(*lhb*:hrGfpII/*fshb*:DsRed2) line to separately target Fsh and Lh cells for electrophysiological experiments. In Fsh cells, we observed oscillatory spontaneous electrical activity (Figure 5A), with cells switching from a non-firing quiescent state to an excited state marked by burst-like sequences lasting 20-80 s.

**Figure 5:**
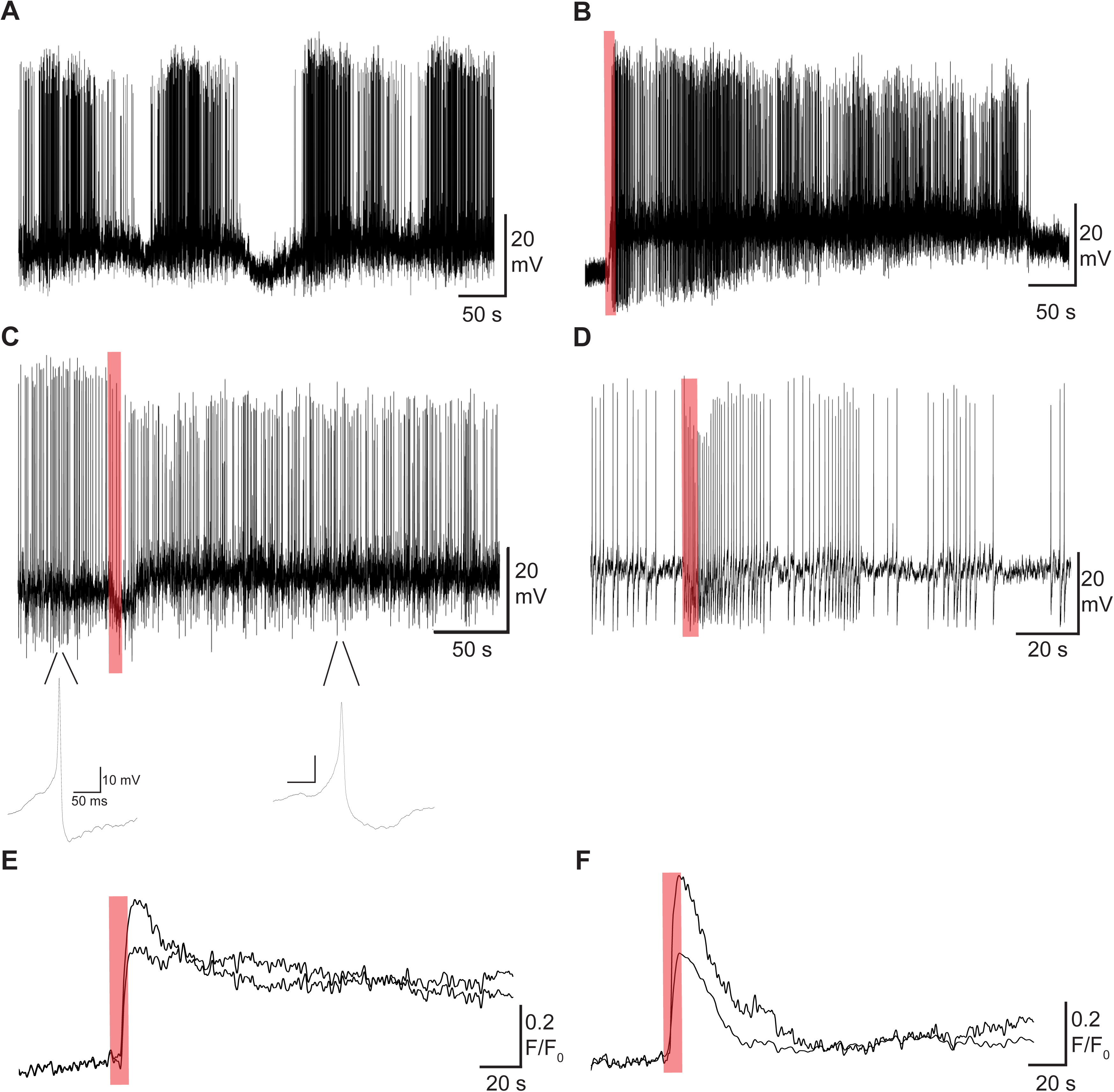
Basal action potential firing properties and Gnrh1-induced responses of Fsh cells in brain-pituitary slices. The experiments were conducted using adult female medaka, tg(*lhb*:hrGfpII/*fshb*:DsRed2) for electrophysiology and tg(*fshb*:DsRed2) for Ca^2+^ imaging. (A) Prolonged current clamp recording of a spontaneous firing Fsh cell. The Fsh cells had oscillatory firing properties where episodes of silence were followed by bursts of action potentials lasting 20-80 s. (B-D) Three types of electrical responses were observed in Fsh cells following Gnrh1 stimulation. (B) Current clamp recording of a non-firing Fsh cell. Gnrh1 application (orange transparent bar overlaying the trace) induced a monophasic response with membrane depolarization to threshold inducing action potentials. (C) Current clamp recording of a spontaneous firing Fsh cell. Gnrh1 elicited a weak biphasic response with an initial hyperpolarization but without any interruption in action potential pattern. The weak hyperpolarization was followed by a weak depolarization and increased action potential duration. (D) Current clamp recording of a spontaneous firing Fsh cell. Gnrh1 elicited a transient increase in firing frequency from 0.5-1 Hz to 2-3 Hz lasting 20-50 s. Two types of Ca^2+^ responses were observed in Fsh cells, prolonged and transient following Gnrh1 stimulation (E and F). (E) Ca^2+^ imaging of an Fsh cell stimulated by Gnrh1. The Fsh cells responded to Gnrh1 with increased [Ca^2+^]_i_ with an initial peak followed by a gradual decline. The responses usually lasted more than 60 s. (F) Transient Ca^2+^ response to Gnrh1 was also observed in Fsh cells, lasting 20-50 s.

Surprisingly, contrary to the results in dispersed cell cultures, about 60% of the Fsh cells in the brain-pituitary slices responded to Gnrh1 (a total of 16 cells from 8 pituitaries) (Figure 5B-F). All responding Fsh cells were located in close proximity to Lh cells. Both electrical and increased [Ca^2+^]_i_ responses were either prolonged (i.e. lasting until the recording ended after 4-6 minutes; Figure 5B, C, and E) or transient (i.e. lasting from 40 s up to 5 minutes; Figure 5D and F) with a 1-5 s latency after addition of Gnrh1. The typical electrical response to Gnrh1 was a membrane depolarization (Figure 5B). In non-firing Fsh cells, the Gnrh1-induced depolarization was sufficient to initiate a prolonged burst. In some cells we observed a weak hyperpolarization prior to depolarization (Figure 5C) without a change in firing activity. In Fsh cells spontaneously generating action potentials, Gnrh1 caused a membrane depolarization and in a few cases led to broadening of the action potential from 6-10 ms to over 20 ms. Some of the Fsh cells also responded to Gnrh1 with small bursts (Figure 5D). The different Gnrh1-induced responses were never observed in control experiments where we applied puff ejections of ECS onto cells. In fact, the electrical activity could not be changed even when doubling the puff application pressure.

The electrophysiological responses of Lh cells in brain-pituitary slices to Gnrh1 were similar to those observed in dissociated cell culture. We saw a clear electrical response in all Lh cells following Gnrh1 stimulation with similar latency as in Fsh cells. In 9 out of 13 cells (from 8 different pituitaries) we observed a biphasic response (Figure 6A). In the remaining 4 Lh cells (3 different pituitaries) out of the 13, we observed a monophasic response in which Gnrh1 initiated a direct depolarization of the cell membrane (Figure 6B). This depolarization was sufficient to initiate both transient and prolonged firing in previously quiescent cells. The electrical responses were also reflected in the different Ca^2+^ responses to Gnrh1 (Figure 6C and D). Prolonged and robust Ca*^2+^* responses to Gnrh1 had two slightly different shapes. One response was clearly biphasic similar to what we observed in cell culture, with an initial Ca^2+^ peak (release of Ca^2+^ from internal stores) followed by a second phase plateau. The other Gnrh1-induced response was prolonged but lack the second plateau following. Following an initial peak, the Ca^2+^ levels gradually decreased but not to basal levels (Figure 6C). In a few cells, we could only detect a transient Ca^2+^ response lasting less than a minute (Figure 6D) before returning to baseline values.

**Figure 6:**
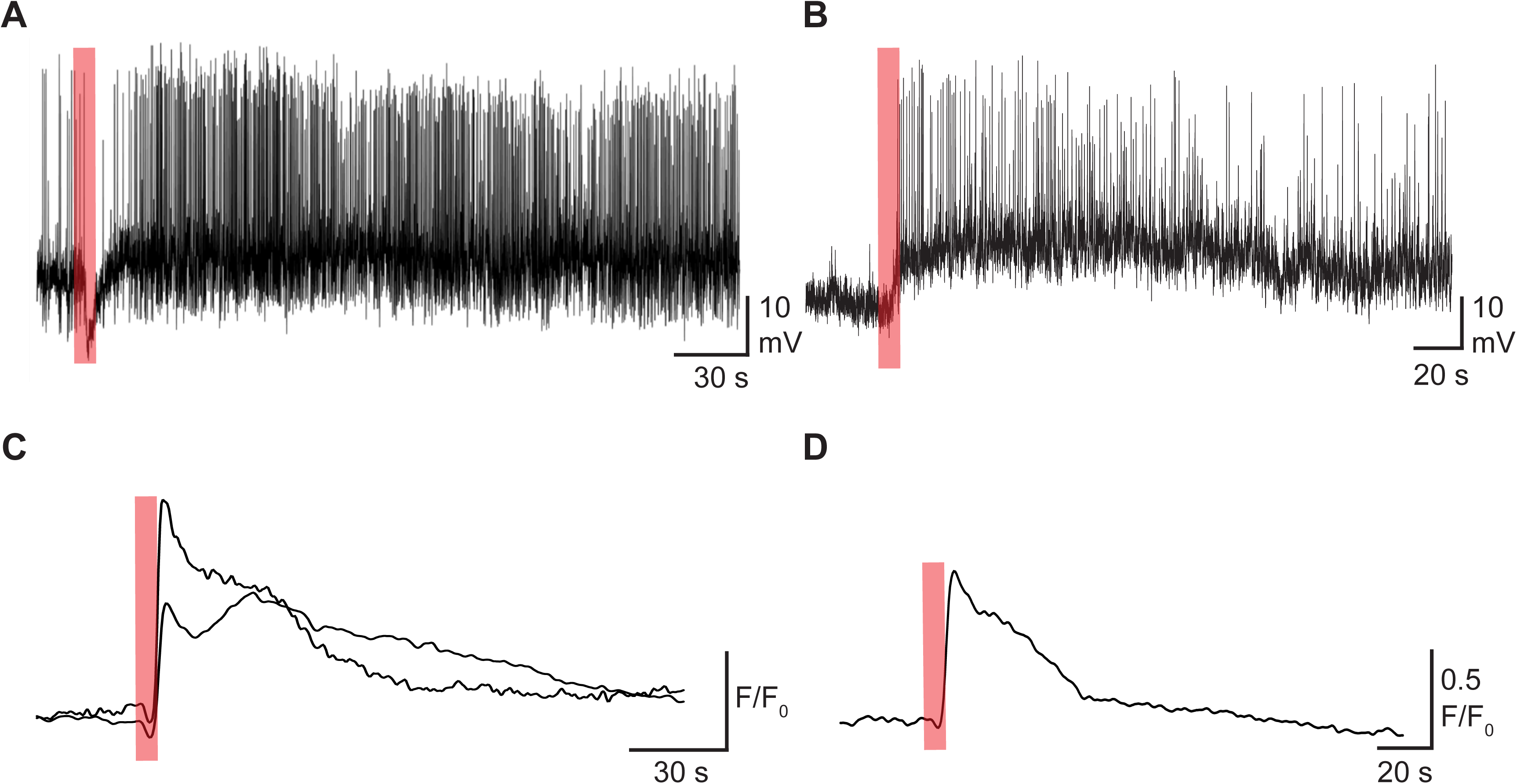
Gnrh1-induced electrical and Ca^2+^ responses of Lh cells in brain-pituitary slices. The experiments were conducted using adult female medaka, tg(*lhb*:hrGfpII/*fshb*:DsRed2) for electrophysiology and the tg(*lhb*:hrGfpII) for Ca^2+^ imaging. Two types of Gnrh1-induced responses could be observed in Lh cells. (A) Typical current clamp recording of a spontaneous firing Lh cell. Gnrh1 stimulation (orange transparent bar overlaying the trace) induced a biphasic response with an initial hyperpolarization followed by a depolarization and increased firing and/or increase action potential duration. (B) Current clamp recording of a quiescent Lh cell. Gnrh1 elicited a monophasic response with a depolarization of the cell membrane to threshold, inducing action potentials. (C) Ca^2+^ responses of an Lh cell stimulated by Gnrh1. The Lh cells responded to Gnrh1 with either a biphasic increase in [Ca^2+^]_i_ mirroring the electrical response in (A) or a prolonged response with an initial [Ca^2+^]_i_ peak followed by a gradual decrease. The responses usually lasted more than 60 s. (D) A few Lh cells responded to Gnrh1 with a transient elevation in [Ca^2+^]_i_ lasting 20-50 s.

### Cell communication between gonadotrope cells in live brain-pituitary tissue slices

The divergent effects of Gnrh1 on Fsh cells in tissue slices and dissociated cell cultures, and the absence of detectable *gnrhr* mRNA in Fsh cells by FISH, led us to hypothesize that intercellular communication may mediate Gnrh1-induced excitation in Fsh cells. To test for intercellular electrical communication, we utilized Ca^2+^ uncaging in one gonadotrope cell and recorded changes in membrane potential in neighboring gonadotropes (Figure 7A). Uncaging of Ca^2+^ causes membrane hyperpolarization via activation of Ca^2+^-activated K^+^ channels (48). If there is direct electrical communication between two cells, altering the membrane potential in one cell should initiate changes in membrane potential in the other.

**Figure 7:**
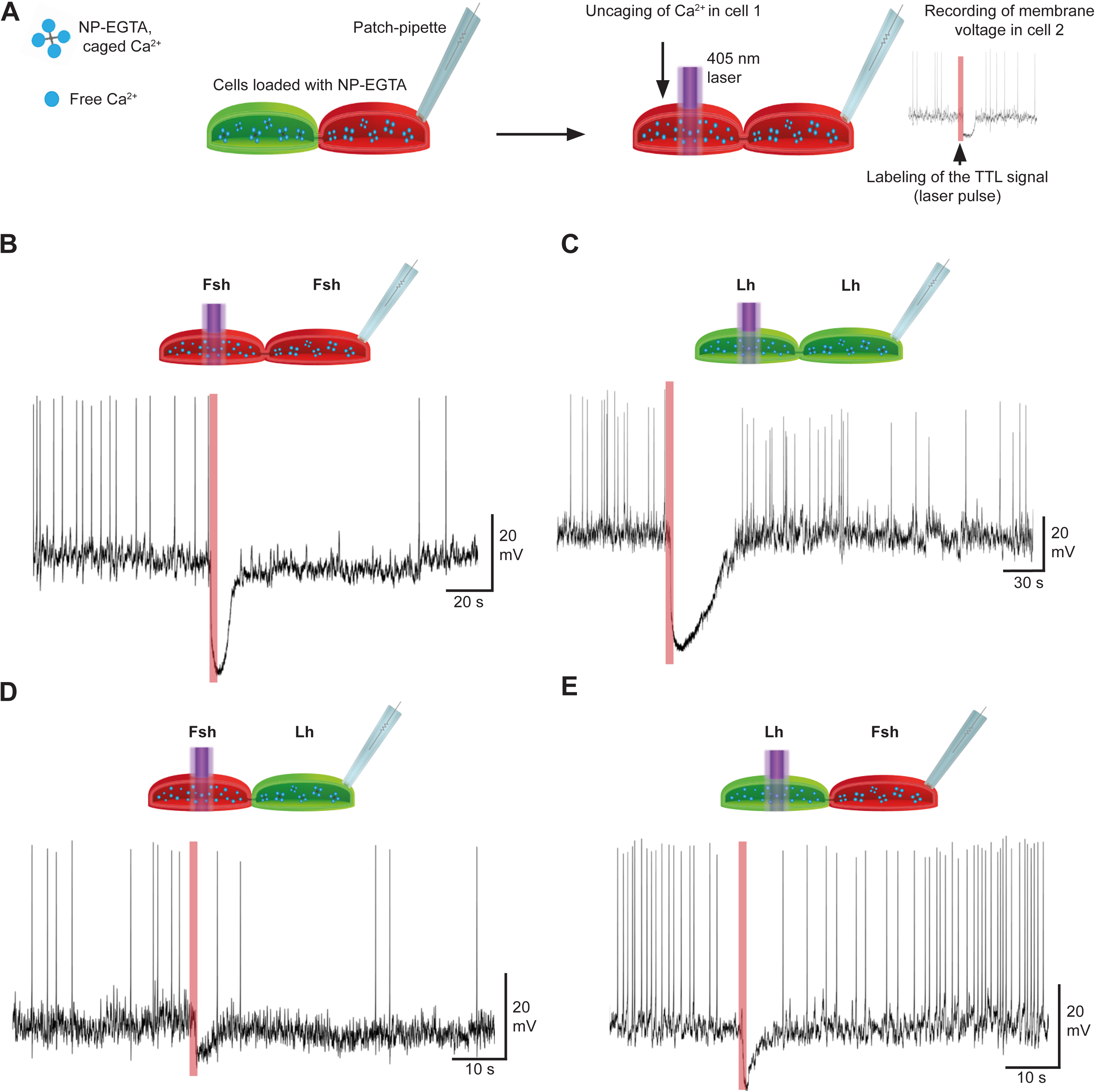
Electrophysiological responses to uncaging of Ca^2+^ in neighboring gonadotrope cells. The experiments were conducted using adult female tg(*lhb*:hrGfpII/*fshb*:DsRed2). (A) A schematic overview of the experimental procedure of simultaneous uncaging and current clamp (voltage) recordings performed on gonadotrope cells from brain-pituitary slices. (B-E) Recording of the electrophysiological response in different gonadotrope cells following uncaging in neighboring cell. (B) Voltage recording of an Fsh cell, uncaging in neighboring Fsh cell. (C) Voltage recording of an Lh cell and uncaging in neighboring Lh cell. (D) Voltage recording of an Lh cell and uncaging in neighboring Fsh cell. (E) Voltage recording of an Fsh cell and uncaging in neighboring Lh cell.

We observed that gonadotropes with soma-soma contact with an uncaged cell hyperpolarized within 15-20 ms of Ca^2+^ uncaging. We saw this rapid response in Fsh cell pairs (n = 11 cells from 3 different pituitaries) (Figure 7B) as well as Lh cell pairs (n = 13 from 3 different pituitaries) (Figure 7C) and more importantly, in Fsh-Lh cell pairs (n = 7 from 3 different pituitaries) (Figure 7D and E). This rapid propagation of the response is to the best of our knowledge too fast for the uncaged Ca^2+^ to diffuse from the target cell to the recorded cell and therefore points to direct electrical connection between the cells. Finally, to further explore intercellular communication, we tested whether Ca^2+^ uncaging could initiate [Ca^2+^]_i_ waves that reached surrounding cells. In fact, uncaging Ca^2+^ in Lh cells could generate small waves that led to elevated [Ca^2+^]_i_ in all Lh cells tested (n = 9 target cells from 3 different pituitaries) (Figure 8A and B. Possible pathways illustrated in 8C). Typically, the Ca^2+^ signal only propagated to neighboring cells reaching maximum of 3 cells from the target cell. In contrast, following uncaging in Fsh cells, we were only able to see propagation of Ca^2+^ signal in 50% of the cells (n = 16 target cells from 3 pituitaries, data not shown).

**Figure 8:**
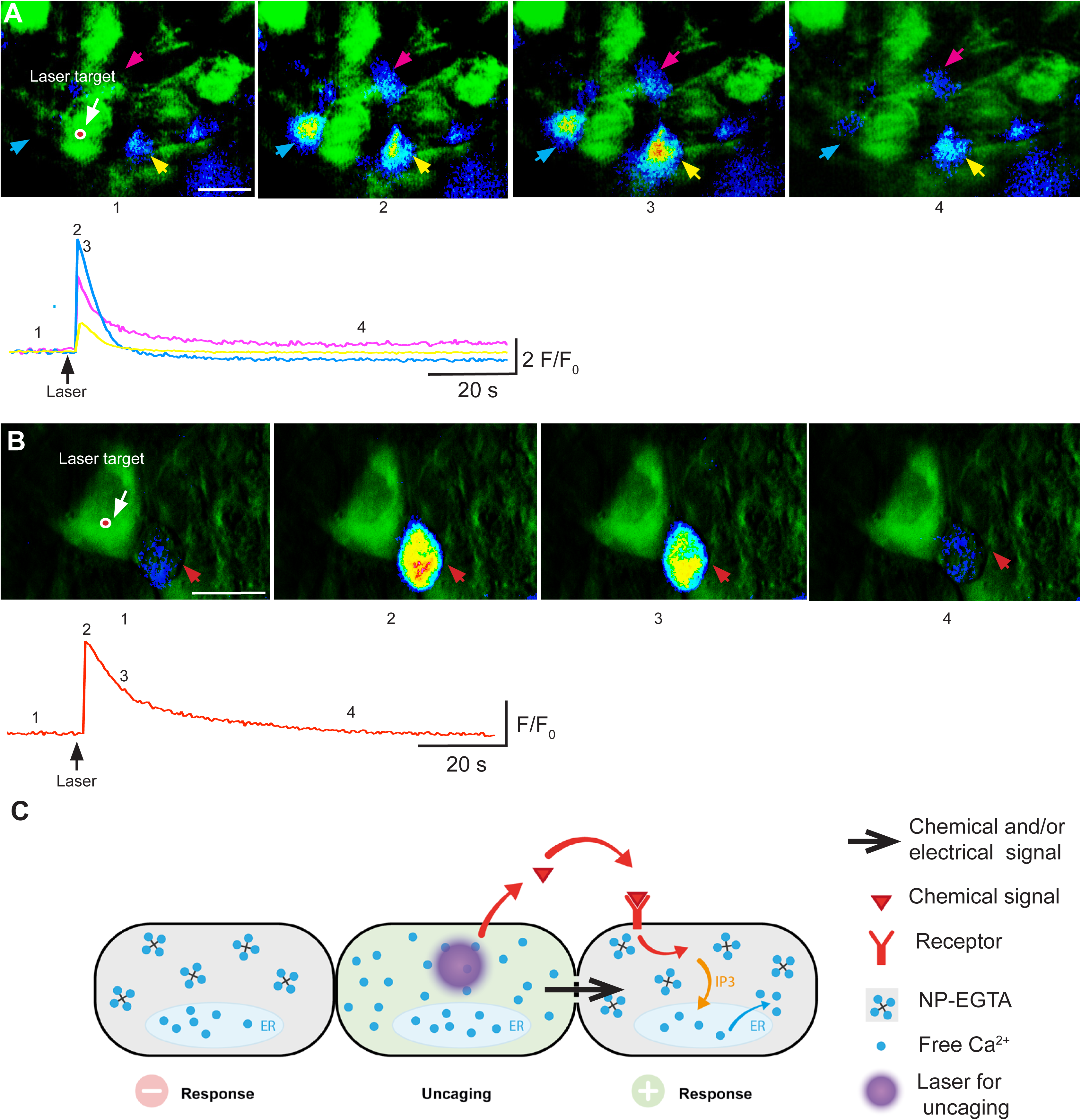
Ca^2+^ responses in surrounding cells following uncaging in Lh cells. (A, B) Ca^2+^ responses in neighboring cells following uncaging in an Lh cell (green color). The laser simultaneously uncaged the Ca^2+^ and bleached the target cell, allowing verification of the precise location of the uncaging. (A) A 300 ms laser pulse targeting one Lh cell elicited a robust Ca^2+^ response in three surrounding cells. Upper panel, representative images of the response. Lower, traces of the three responses calculated as F/F_0_ following background subtractions. (B) Same as (A) but only one neighbor cell responded to the uncaging. (C) Overview of possible mechanisms responsible for the electrical response and the Ca^2+^ response to uncaging of Ca^2+^. Scale bars in A and B: 10 μm.

## DISCUSSION

In this study, we first show that the anatomical organization of the hypophysiotropic Gnrh (Gnrh1 in medaka) is similar to other teleost species (2–5). The Gnrh1 axons projects close to both Fsh and Lh cell as well as the pituitary blood vessels. Following, we find evidence that in female medaka, Gnrh1 stimulates Lh cells directly but Fsh cells indirectly, likely via interactions with directly activated Lh cells. This evidence is three-fold: 1) Fsh cells in female medaka lack expression of any of the three *gnrhr* paralogs present in the pituitary, whereas all Lh cells express at least one *gnrhr* paralog. 2) While Lh cells exhibited identical electrical and Ca^2+^ responses to Gnrh1 in dissociated pituitary cultures and in brain-pituitary tissue slices, Fsh cells showed no effect in dissociated pituitary cultures but did respond to Gnrh1 in brain-pituitary tissue slices. 3) Direct Lh cell activation rapidly induced electrical responses in neighboring Fsh cells. In addition, uncaging of Ca^2+^ in gonadotropes can initiate small [Ca^2+^]_i_ waves that propagate to surrounding cells.

### Organization of gonadotrope cells and Gnrh1 fibers within the pituitary

To separately investigate Fsh and Lh cells we established a new tg line with expression of red fluorescent protein, Dsred2, controlled by the endogenous medaka *fshb* promoter. We confirm the specificity of *dsred2* in *fshb* cells with multicolor FISH. Moreover, Burow et al (40) reported that *fshb* cells also express Fshβ protein in the medaka pituitary. Thus, we inferred that the *dsred2*-positive cells produce Fsh hormone and can therefore be referred to as Fsh cells. We then crossed tg(*fshb*:DsRed2) fish with fish from the previously established tg(*lhb*:hrGfpII) line (36, 47) (Figure 1E-H) and examined the spatial distribution of Fsh and Lh cells. In agreement with prior studies in teleosts (7), in the double tg(*lhb*:hrGfpII/*fshb*:DsRed2) medaka Fsh and Lh were expressed by distinct pituitary cells, thereby validating its use to simultaneously study Fsh and Lh cells (Figure 1I-L).

Lh cells were found to be clustered in the ventral and lateral surface of the pituitary, whereas Fsh cells appeared more spread out and generally located more dorsally than Lh cells. However, Fsh cells were often found in close proximity, and in some cases appeared in direct contact, with other Fsh cells. Significantly, Fsh cells also appeared to make direct contact with Lh cells along the ventral line of the pituitary. Similar observations have been made in zebrafish (*Danio rerio*) where Fsh cells were found at the periphery of Lh-cell clusters (49).

Similar to that reported for other fish species (2–5), we found that Gnrh1 neurons directly innervate the pituitary in female medaka. Gnrh1 projections were seen throughout the PPD where both Fsh and Lh cells are located, and alongside blood vessels (Figure 2A-J). This organization is quite similar to that of Gnrh3 neurons in zebrafish (5). We did not see projections terminating directly on gonadotrope cells, and therefore cannot determine if Fsh and Lh cells are directly targeted. Golan et al (5) reported that Gnrh3 neurons innervating the pituitary in adult zebrafish had varicosity-like structures or boutons. In female medaka, we could not find such structures, but because we did not label the Gnrh1 neurons with synaptic markers, we cannot rule out their existence. We did detect small blebs in close contact with blood vessels (Figure 2K and L). However, during pituitary sectioning, the orientation of blood vessels and Gnrh1 neurons relative to the direction of the cut may introduce bouton-like artifacts in transversally cut neurons. Therefore, additional studies are needed to clarify the exact morphology of the Gnrh1 projections and the precise location of the terminals.

### Lh cells express *gnrhr*, but Fsh cells do not

Four *gnrhr* paralogs have been identified in the medaka genome and expressed in the adult medaka brain (50). In a previous study, two of the *gnrhr* paralogs were found to be expressed in the adult female medaka pituitary, namely *gnrhr1b* and *gnrhr2a* (21). In the present study, we detected one additional paralog; *gnrhr2b*. Expression of multiple *gnrhr* paralogs in the pituitary has been observed in other teleost species including goldfish (*Carassius auratus*)(51), tilapia (*Oreochromis niloticus*)(28, 52) and (*Astatotilapia burtoni*)(53), and Atlantic cod (*Gadus morhua*)(54). Using *in situ* hybridization on brain-pituitary slices from female medaka, we found *gnrhr* expressed in the median and posterior part of the pituitary. While *gnrhr1b* and *gnrhr2b* were expressed in the PI, only *gnrhr2a* was expressed in the PPD, and almost exclusively in Lh cells (Figure 3). These results support a previous study where *gnrhr2a* was the paralog showing the highest expression levels in the pituitary of both juvenile and adult medaka (50). The same study revealed that *gnrhr2a* expression levels increased in parallel with the number of pituitary Lh cells between juvenile and adult fish (50).

Interestingly, *lhb* cells from Atlantic cod were found to express *gnrhr1b* and *gnrhr2a* (54). More recently, a novel *gnrhr*, *gnrhr2baα*, was found in *lhb* cells from Atlantic salmon (29). It is interesting to note that the *gnrhr* identified in Lh cells from Atlantic salmon, Atlantic cod, and medaka belong to the same phylogenetic group, suggesting they share a common ancestral gene (29).

To our surprise, we did not co-localize any *gnrhr* with *fshb*. This contradicts previous reports of *gnrhr* expression in *fshb* cells from Atlantic cod (19, 54) and tilapia (28). This disparity may be due to species or sex differences, or to differences in methodology. The work in Atlantic cod was performed on primary dissociated pituitary cells pooled from both sexes and conducted at 2 to 7 days after plating (19, 54). This difference in timing may be important, as a previous study found that *gnrhr* expression increased in pituitary cell culture over time (55), suggesting that either *gnrhr* expression increases in cells already expressing the receptor or that new cells start to express *gnrhr*. In tilapia, single-cell PCR found measurable *gnrhr* transcripts in *fshb* cells in fixed tissue; however, this study only analyzed pituitaries from males (28). Therefore, further studies are needed to clarify whether *gnrhr* expression patterns vary among species or between sexes. Notably, the expression pattern we report for female medaka is similar to that observed during embryogenesis in mouse, where FSH and LH are produced in distinct cells within the pituitary and during this time GnRHR are expressed exclusively in LH cells and not in FSH cells (56).

### Effects of Gnrh1 on Fsh and Lh cells in cell culture

In the present study both the electrophysiological recordings and Ca^2+^ changes in cultured cells treated with Gnrh1 are consistent with the *in situ* hybridization results. We found that female medaka Lh cells clearly responded to Gnrh1, inducing elevated [Ca^2+^]_i_ and altered electrophysiological behavior. Conversely, we did not detect changes in membrane potential or Ca^2+^ levels in Fsh cells upon exposure to Gnrh1 up to 48 h after plating the dispersed pituitary cells. These results confirm that the *in situ* hybridization is adequately sensitive and all together suggest that unlike Lh cells, adult female medaka Fsh cells do not express *gnrhr*.

The response of Lh cells to Gnrh1 is consistent with that previously reported for other teleosts (19,21,22,43). However, very few studies have examined electrical activity or calcium flux of cultured teleost Fsh cells in response to Gnrh1. In dispersed Atlantic cod pituitary cultures, Gnrh increased action potential frequency and [Ca^2+^]_i_ in Fsh cells, suggesting that Fsh cells possess functional Gnrhr (19). However, as noted above, that study was conducted after the dispersed pituitary cells had been maintained for 2 to 7 days in culture, which may have induced phenotypic changes such as altered *gnrhr* expression. As a result of these contradictory results, a more systematic testing of Gnrh induced responses using primary pituitary cells should be conducted to reveal if gonadotrope cells alter their Gnrhr composition with time in culture.

### Effects of Gnrh1 on Fsh and Lh cells in brain-pituitary tissue slices

Both Fsh and Lh cells fired spontaneous action potentials in brain-pituitary slices (Figure 5 and 6). In addition, long current clamp recordings of Fsh cells demonstrated membrane potential oscillations, with quiescent periods followed by weak depolarization and subsequent firing of action potentials. These longer bursts of spontaneous action potentials could mediate basal release of Fsh, but further experiments are needed to resolve their exact role. Using inverse-Pericam transgenic medaka, Karigo et al (20) observed that unstimulated Lh cells exhibited regular synchronized [Ca^2+^]_i_ oscillations. Fsh cells had more desynchronized [Ca^2+^]_i_ oscillations with lower peaks compared to Lh cells. However, the frequency of [Ca^2+^]_i_ peaks were higher in Fsh cells. Importantly, the [Ca^2+^]_i_ oscillations were demonstrated to be independent of Gnrh (20).

Surprisingly, despite finding no expression of *gnrhr* in Fsh cells, and no response to Gnrh1 in dissociated Fsh cells, Fsh cells in brain-pituitary slices did respond to Gnrh1 (Figure 5). This result agrees with findings reported by Karigo et al (20) who showed that Gnrh1 elicited elevated [Ca^2+^]_i_ levels in Fsh cells in whole brain-pituitary preparations from female medaka (20). Furthermore, this is the first clear demonstration that the effects of Gnrh1 on Fsh cells are indirect.

### Cellular communication between gonadotrope cells

Because Fsh cells respond to Gnrh1 in tissue slice preparations but not in dispersed cell cultures, and because they lack functional Gnrh receptors, we explored the possibility of communication between gonadotrope cells in pituitary slices (Figure 8). By using Ca^2+^ uncaging and patch clamping on genetically labeled Fsh and Lh cells, we separately targeted the cells within the same preparation. Indeed, current clamp recordings demonstrated that not only is homotypic communication between Fsh cells or between Lh cells possible, but Fsh and Lh cells form heterotypic networks as well. Such cell-cell networks allow rapid electrical communication, whereby information received by one cell can be quickly relayed to several cells. Coupling of cells in the pituitary has been shown to greatly affect hormone release in both mammals and teleost fish (33,34,57,58). However, blocking gap junctions responsible for cytosolic bridging between cells had less impact on Gnrh induced Fsh release than Lh release in tilapia (34). This observation is consistent with our finding that the responses of Fsh cells to Gnrh1 were weaker than that of Lh cells. In fact, as previously reported, normal folliculogenesis is seen in Gnrh knockout medaka (13). In addition, in mammals only a 50% reduction in plasma FSH is observed in GnRH deficient *hpg* mouse (30). These results suggest that FSH is more dependent on other factors than GnRH. Therefore, additional investigations are required to identify all factors responsible in the differential regulation of Fsh and Lh.

In our experiments we clearly see that Ca^2+^ uncaging in single cells can propagate between Lh cells (Figure 8) which may explain the synchronicity of the Ca^2+^ flux in Lh cells observed by Karigo et al (20). Future studies combining uncaging with Ca^2+^ imaging could reveal potential differences between Fsh and Lh cells in how Ca^2+^ propagates between the cells.

To conclude, we provide evidence that while Lh cells in female medaka respond directly to Gnrh1, Fsh cells do not. However, Fsh cells can respond to Gnrh1 when they are associated with other pituitary cells. We also provide evidence of electrical signaling among gonadotropes. We propose that such signaling may play an important role in gonadotrope physiology and suggest that intercellular electrical signaling may mediate hormone release and permit Fsh cell response to Gnrh1.

## ACKNOWLEDGEMENTS

We thank Drs Susann Burow and Rasoul Nourizadeh-Lillabadi for help with development of the medaka transgenic line, tg(*fshb*:DsRed2), and Dr Felix Loosli (Karlsruhe Institute of Technology, Germany) for kindly providing the I-SceI-MCS-leader-Gfp-trailer plasmid.

